# The spatial layout of antagonistic brain regions is explicable based on geometric principles

**DOI:** 10.1101/2024.06.17.599309

**Authors:** Robert Leech, Rodrigo M Braga, David Haydock, Nicholas Vowles, Elizabeth Jefferies, Boris Bernhardt, Federico Turkheimer, Francesco Alberti, Daniel Margulies, Oliver Sherwood, Emily JH Jones, Jonathan Smallwood, František Váša

## Abstract

Brain activity emerges in a dynamic landscape of regional increases and decreases that span the cortex. Increases in activity during a cognitive task are often assumed to reflect the processing of task-relevant information, while reductions can be interpreted as suppression of irrelevant activity to facilitate task goals. Here, we explore the relationship between task-induced increases and decreases in activity from a geometric perspective. Using a technique known as kriging, developed in earth sciences, we examined whether the spatial organisation of brain regions showing positive activity could be predicted based on the spatial layout of regions showing activity decreases (and vice versa). Consistent with this hypothesis we established the spatial distribution of regions showing reductions in activity could predict (i) regions showing task-relevant increases in activity in both groups of humans and single individuals; (ii) patterns of neural activity captured by calcium imaging in mice; and, (iii) showed a high degree of generalisability across task contexts. Our analysis, therefore, establishes that antagonistic relationships between brain regions are topographically determined, a spatial analog for the well documented anti-correlation between brain systems over time.

**Significance Statement:** It is well documented that brain activity changes in response to the demands of different situations, although what gives rise to the observed cortical activity patterns remains poorly understood. Using analytic tools from earth sciences, we examined whether the landscape of regional changes in activity emerge from a set of common topographical causes. Using only regions showing decreases in activity, we could predict the landscape of regions showing increases in activity using fMRI in humans and calcium imaging in mice. Our results suggest topographical principles determine the landscape of peaks and valleys in brain activity -- a spatial analog for the well documented anti-correlation between sets of brain regions over time.

## Introduction

One of the most well-established features of brain activity is a landscape of regional increases and decreases in activity that emerge in humans and other mammals. For example, a distributed set of regions show systematic decreases in activity when participants engage in a range of complex tasks^1^. These regions are known as the default mode network, and have been hypothesised to play a role in cognition and behaviour linked to memory and other internally focused cognitive processes^2^. At the same time, a different set of regions show a tendency to increase activity when participants perform tasks with increased cognitive demands, a system which is often referred to as the multiple demand network and is assumed to play an important role in executive control^3^.

Functional accounts typically argue that task-positive or task-negative features of brain activity reflect how the brain implements different features of cognition, such as the application of task rules in the case of executive control^3,4^ or the use of long-term knowledge to guide cognition and behaviour in the case of the default mode network^2^. In contrast, in our current study, we consider a complementary hypothesis: that regional increases and decreases in brain activity observed during different states are the result of a set of common topographic causes. To investigate this possibility, we examined whether the spatial distribution of regions that show increases in brain activity can be predicted by the regional distribution of regions that show reductions in brain activity (and vice versa).

The importance of physical space as an organising principle has long been recognised as fundamental for brain function. For instance, sensory and motor functions are arranged as topological maps^5^ and processing streams^6,7^. However, an increasing number of studies^2,8–12^ have highlighted the possibility that geometric constraints may provide a set of general principles which could explain the rich neural dynamics that are observed across the cerebral cortex and would therefore help constrain how these neural processes support different cognitive functions.

These geometric perspectives suggest that the specific spatial patterns of activation and deactivation we observe with neuroimaging reflect the operation of topographical constraints that relate to how the brain functions as a system. An analogy can be made with landscapes, where the pattern of elevation on the Earth’s surface reflects processes that are distributed in space (e.g., plate tectonics, volcanic activity, glaciation, and weathering). In many of these phenomena the peaks and valleys observed within a specific landscape are a consequence of a small number of geological processes which lead to predictable changes in elevation over space and time that are common across landscapes located in different parts of the globe. Extrapolating from this evidence to brain function, we hypothesised that the relative increases and decreases in brain activity that emerge during specific task states may be in part determined by the operation of a finite number of topographical principles that constrain brain activity on the cortex. Such a view of brain function would be supported by evidence that the pattern of regional increases in activity can be predicted by the distribution of regions that show decreases in activity (and vice versa).

In order to determine whether common topographical principles explain both increases and decreases in brain activity during tasks we used an approach that is commonly used to infer topographical features in geology and other earth sciences. It is well established that because processes like glaciation shape both the peaks of mountains and valley floors. Consequently, physical descriptions of the shape of a mountain peak contains information that also describe the shape of the valley floor (and vice versa). In a geological context, the relationship between peaks in valleys can be captured by a spatial regression technique called Kriging^13^ that has been applied in other domains such as ecology, geography, and climate sciences. Kriging has recently been extended to work with large datasets of tens of thousands of observations and accommodate spatial heterogeneity (i.e. the possibility that underlying influences on a landscapes topography can vary across space), making it a viable technique for facilitating its use in a neuroimaging context^14^.

In our study we apply Kriging (Figure 1, top) to the distribution of brain activity observed in a range of different situations and using different imaging techniques (fMRI and calcium imaging). Specifically, we examined whether the spatial distribution of vertices that show reductions in activity (i.e., vertices on the cortical surface that show a relative deactivation) are able to predict the distribution of vertices that show activity increases (i.e., vertices with a relative increase in activity) and vice versa (Figure 1, bottom). Our findings highlight that this can be done with remarkable accuracy indicating that: (i) many of the observable features of task-positive brain activity are spatially linked to reductions in activity (and vice versa); (ii) that this is not tied to a specific modality of brain imaging and (iii) while there are unique task-positive activity patterns arising from the spatial configuration of task deactivation for each task there are also important similarities seen when human perform different cognitive tasks, explaining why often different tasks can have similar neural profiles. Together these analyses establish that the task-positive and task-negative dimension of brain activity are at least partly an emergent property of cortical geometry. More generally, they underscore the importance of mapping both regional increases and decreases in activity in theoretical approaches to the brain basis of cognition.

**Figure 1:**
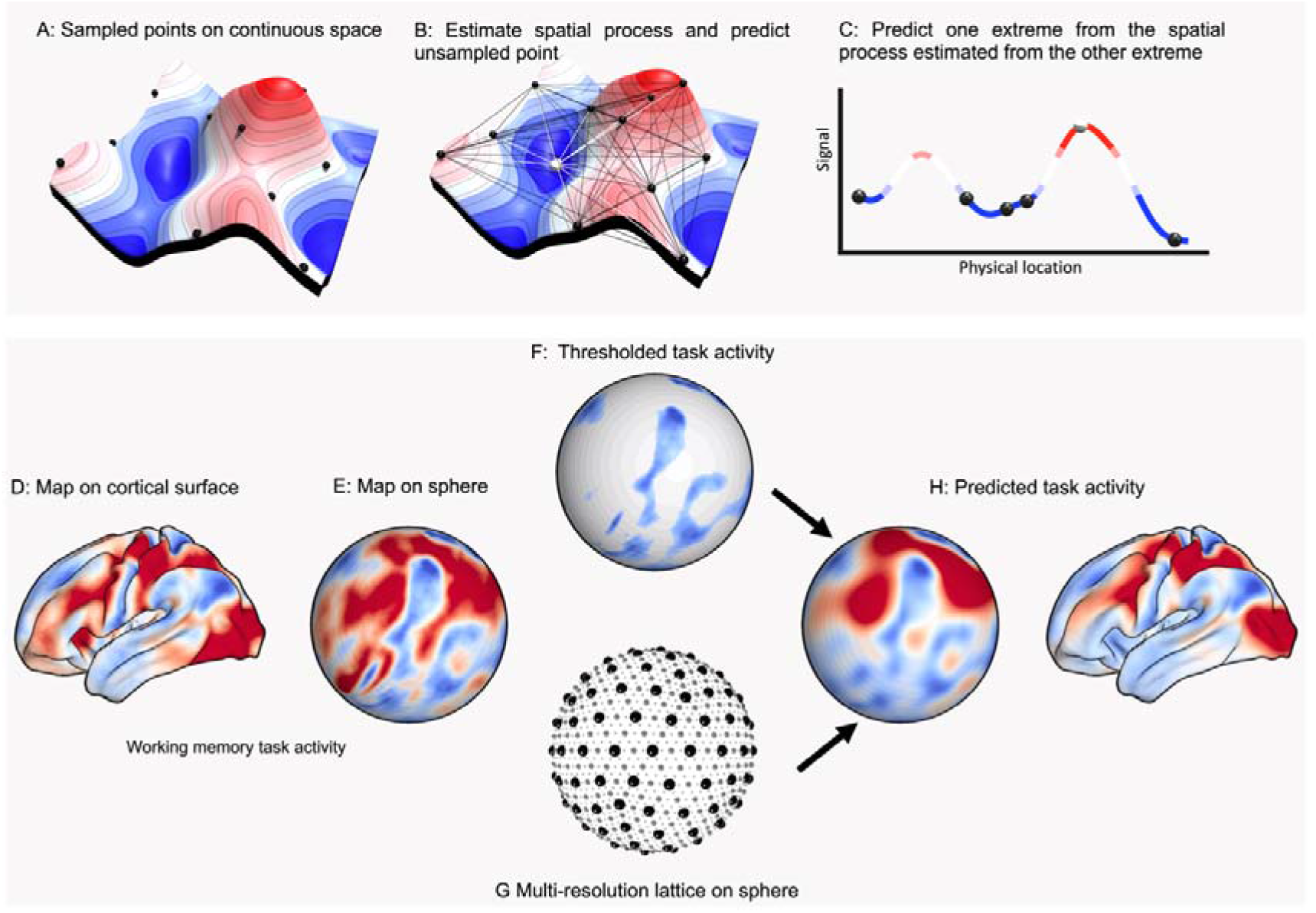
The spatial regression approach (Kriging). Top, schematic of the spatial regression approach: A) an illustration of a signal (elevation and color) in a 2D physical space, with black spheres indicating sampled points where the signal has been measured.B) Distances and similarities of the signal between sampled points (black lines) can be used to approximate the spatial process of the continuous signal. This spatial process can be used to predict unsampled points (white sphere) based only on distance to sampled points (white lines). C) 2D cut through, showing using samples (black points) from one extreme to predict the other extreme (white point). Bottom, application of the spatial regression approach to the cortical surface. D) A cortical surface map is identified; for example, activations from a cognitive task. E) The map is projected onto a sphere, and F) thresholded to retain a subset of regions; here, those that show deactivation for the chosen task. Next,G) a multiresolution regular lattice is created on the sphere, to estimate spatial covariance and H) perform spatial prediction of the whole-brain map, including in particular the out-of-sample (task-positive) regions. Finally, these can be compared to the true out-of-sample values (i.e., observed task-positive activity).

## Results

We first demonstrate the utility of kriging as a technique for explaining the spatial distribution of increases and decreases in brain activity using group average BOLD activity from the Human Connectome Project^15^. We took a single group-averaged contrast z-score map (e.g., 0-Back relative to baseline) for each of the seven cognitive tasks (see Figure 2). Each task map was thresholded to retain the lowest 25% of cortical vertices (i.e., most negative values; excluding the medial wall) and the resulting mask used as the input for spatial prediction. We performed lattice Kriging by projecting the masked input data onto a multiresolution spherical lattice and using the pattern of changes in activity as a function of distance to approximate the underlying spatial process and based on this, predict all vertices on the cortical surface (except the medial wall). The model was estimated separately for each task, without prior training on other datasets; as such, each separate model was only provided with the thresholded spatial distribution of activity decreases for that specific task. In this analysis predictions are entirely based on regions showing decreases in activity and we assess whether this is able to predict the distribution of positive activity in the unseen vertices. Consistent with our hypothesis that a set of common topographical principles explains both increases and decreases in brain activity during tasks we (Figure 2A) generated a set of predictions for each task (Figure 2B), which showed reasonable correspondence to the ‘true’ task patterns (Figure 2C) (see Supplementary Figure 1 for the inverse predictions, i.e., predicting task negative vertices from super-threshold task-positive vertices).

**Figure 2:**
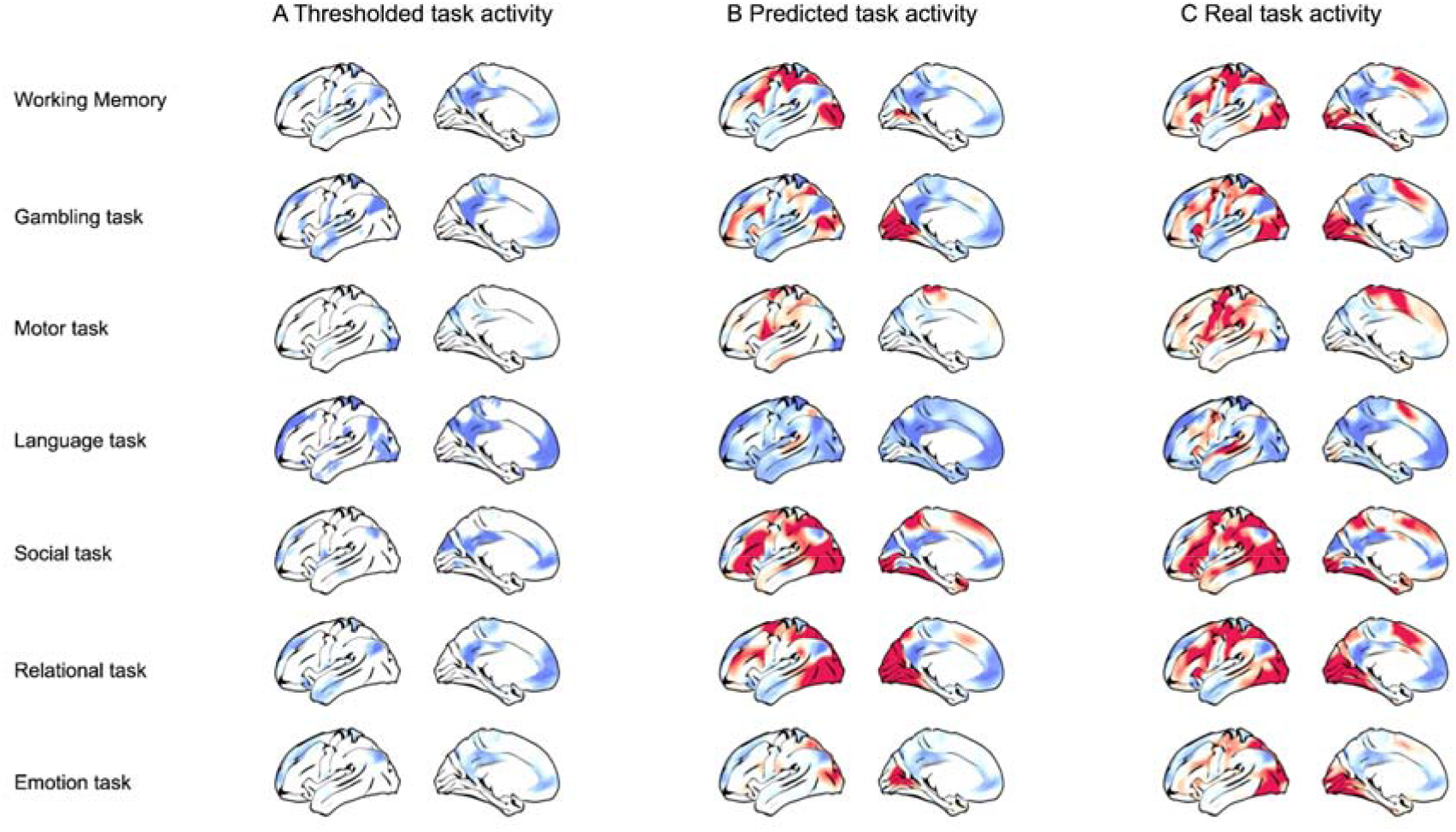
Spatial prediction from task-negative activity patterns. **A**) Vertices with the lowest (< 25%) group-average task activation were used to form a predictor mask. **B**) Spatial regression was used to predict task activity at all remaining vertices, and compared to **C)** the true pattern of activity.

We quantified the relationship between predicted and true increases in brain activity in the out-of-sample vertices (Figure 3A). Spearman’s correlations between predicted and empirical activity ranged from *ρ* = 0.57 (for the Emotion task) to *ρ* =0.77 (for the Theory of Mind task). The presence of autocorrelation unrelated to task activity (e.g., resulting from thermal noise, registration error between subjects, etc.) means that some positive correlation in out-of-sample prediction is to be expected by chance. Therefore, we also calculated the correlation between true and predicted activity within a restricted set of vertices (Figure 3B). We restricted the correlation between real and predicted activity to vertices with task-positive activity (i.e., z-score>0), excluding any vertices outside of the predictor mask whose activity was negative (z<0). This restricted analysis still showed positive correlations between real and predicted activity (range of Spearman’s *ρ* across tasks = 0.34-0.8); demonstrating that the model does not purely predict the *location* of vertices with positive BOLD activity, but also the *variability* in activity across these task-positive vertices.

**Figure 3:**
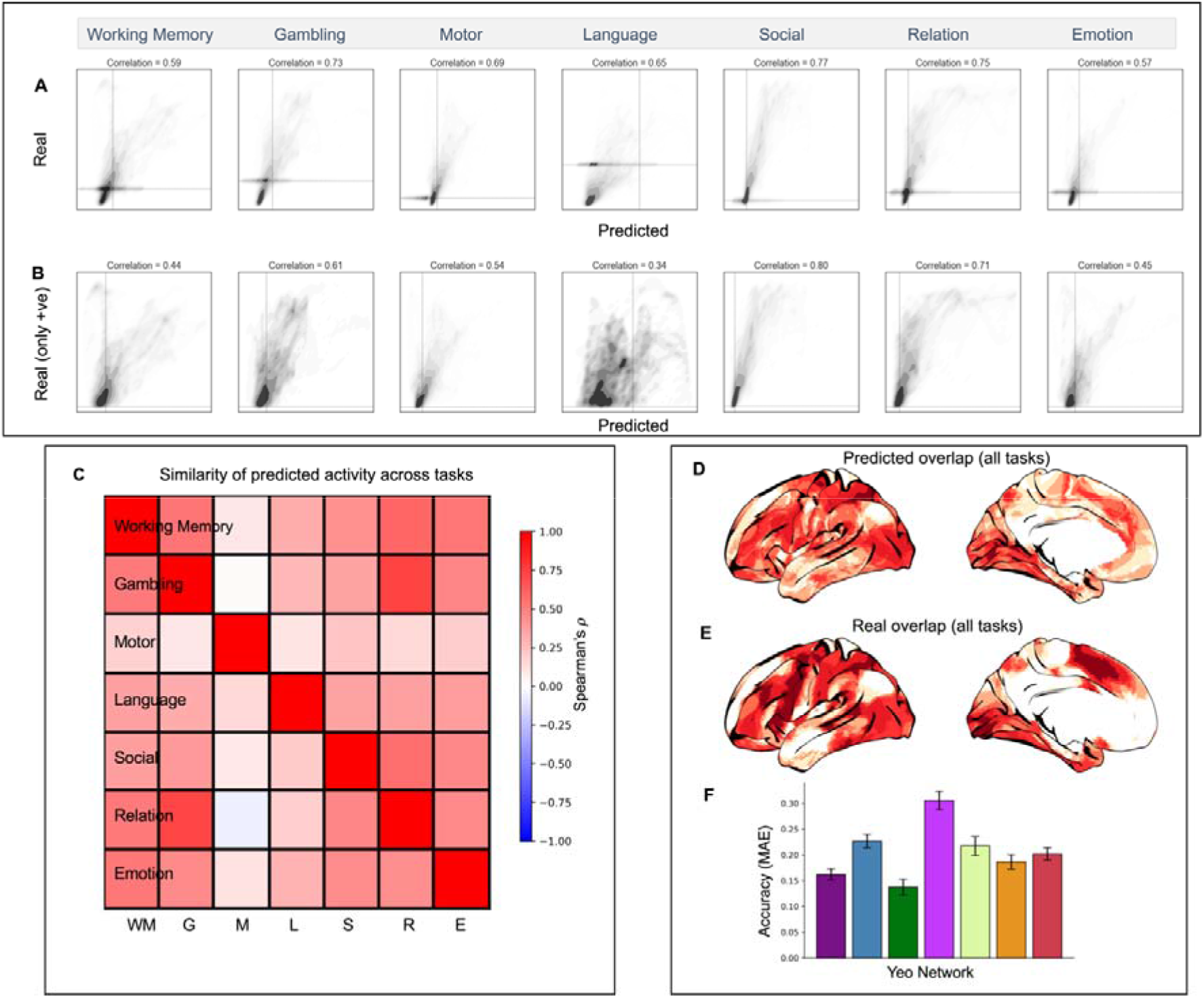
Assessing the agreement between predicted and true patterns of activity. Left panel, **A**) scatter plots of the relationship between true and predicted out-of-sample vertices for each task. **B**) The relationship between predicted and real activity restricted only to vertices with a true task-positive BOLD response (i.e., z>0). **C)** Similarity across tasks of predicted activity for all out of sample vertices. **D**) The overlap of positive predicted values across all tasks; **E**) the real overlap of positive values across all tasks; and **F**) accuracy (mean absolute error) for predicted overlap (of out-of-sample positive values) for different intrinsic connectivity networks (Yeo*, Krienen* et al, 2011); the mean and standard deviation of the accuracy was calculated by leave-one-task out (see methods for details).

In Figure 3C, we also observe the similarity in out-of-sample predictive performance (derived from each task-negative pattern of activity) across each of the 7 tasks. This illustrates that for the majority of the tasks (with the exception of the motor task), the predicted solution is well-aligned across tasks (although it is always strongest for prediction of its own out-of-sample activity). This indicates a mixture of task-general and task-specific aspects of the task-negative spatial configuration. An informative summary measure is the overlap in the number of tasks predicted to have positive activity at each vertex (Figure 3D-E), regions that tend to show good spatial agreement between predicted and real data. Figure 3F shows the predictive accuracy of the overlap measure in 3D-E stratified by intrinsic connectivity networks^16^; the worst prediction (highest mean absolute error; MAE) was found for the ventral attentional network, while the best prediction (lowest MAE) was found for the dorsal attention network.

In Figure 3D-F, we observed evidence of both a shared pattern of task-positive activity predictable from task negative activity for the majority of tasks as well as elements that are unique to each task.

To further evaluate the task-specific component of the spatial prediction, we next directly compared predictive performance across pairs of tasks (Figure 4, top). We generated a binary mask based on the intersection of a pair of (previously-generated) task-specific masks. We used Kriging to reconstruct the underlying cortical activation maps from a subset of task activity which was spatially restricted only to the intersection mask (i.e. we predicted both increase and decreases in brain activity). Finally, we compared the predicted map to both tested empirical task maps, to quantify whether the predicted map is more similar to the corresponding empirical map (whose activation within the intersection mask was used for prediction). We repeated this across all 42 pairs of tasks. This allowed us to systematically compare predictive performance for pairs of tasks based on matching subsets of vertices in the predictor mask. As such, this was a highly constrained test of the hypothesis that the spatial distribution of task activity within the predictor mask allows for meaningful spatial prediction outside of the mask. Figure 4, bottom, shows the comparison of out-of-sample similarity between each pair of tasks (true versus alternative task). Out of all 42 task comparisons, the true task used to generate the prediction had greater predictive performance than the alternative task 39 times (Figure 4, bottom A). Similar results were obtained when the out-of-sample performance was restricted to either a minimum distance from any vertex in the mask (0.1 radians, 3 mm; Figure 4, bottom B), or only to vertices that had positive activity in the real pattern of activity (Figure 4, bottom C).

**Figure 4:**
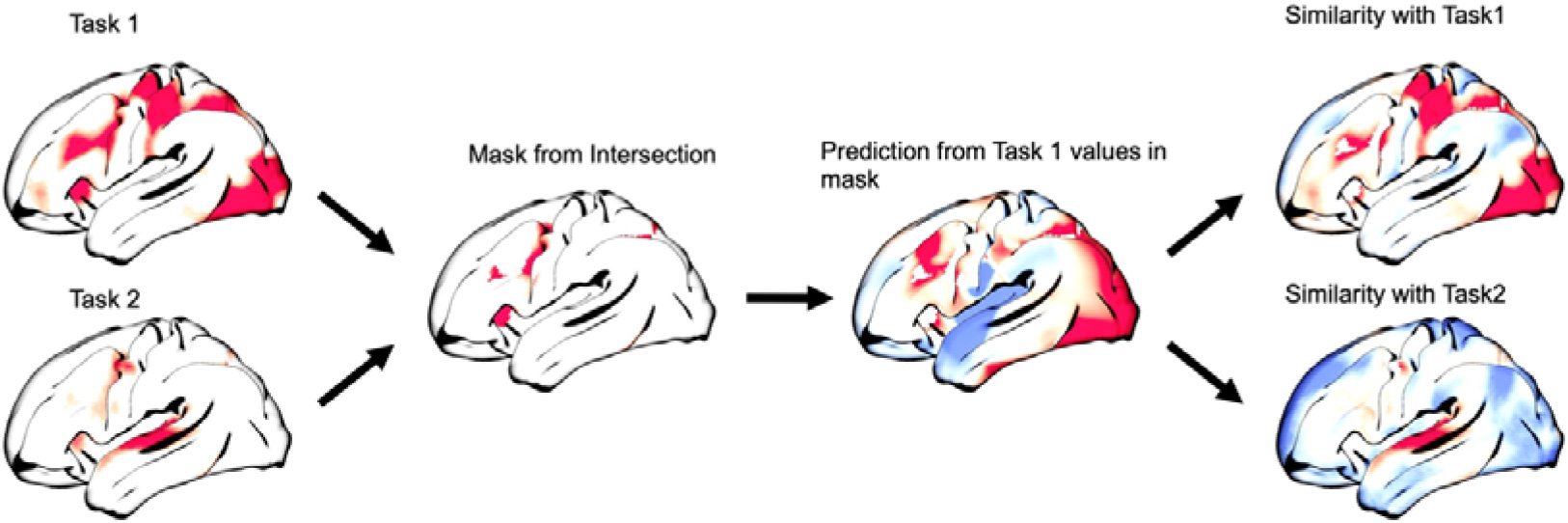
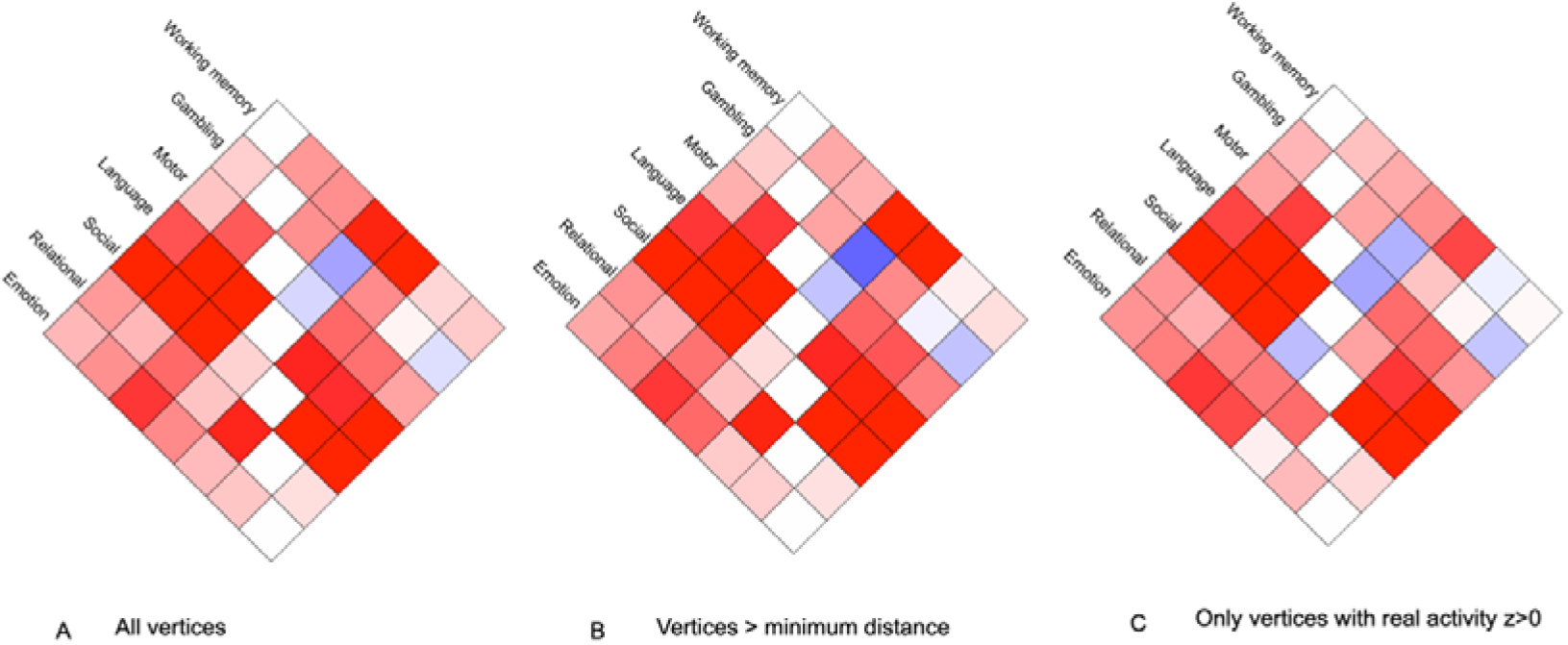
Assessing the task-specificity of spatial predictions. Top: The approach consists of creating a mask from the intersection of two task-specific masks, predicting activation for one task across the rest of the cortex, and then assessing whether the predicted output is more similar to the (expected) true task than to the alternative task. Bottom: A, Pairwise similarity in true versus alternative task performance, based on: A) all out-of-sample vertices;B) out-of-sample vertices further than 0.1 radians from any vertex in the predictor mask; C) out-of-sample vertices with empirical activity z>0.

So far, we have shown that the input masks based on regional decreases can predict regions that show positive activity. A complementary perspective on spatial prediction can be obtained by constraining input data to predefined brain regions or networks, and assessing their relative ability to predict activity in the rest of the brain. Therefore, we also defined masks from seven canonical intrinsic connectivity networks^16^ and used these to generate spatial predictions of activations for vertices outside of each mask. Figure 5 shows the correlation between out-of-sample real and predicted activity for each network and each task. Across tasks, the networks covering large primary sensorimotor systems (i.e., visual and somatomotor networks) performed poorly; conversely, association networks (fronto-parietal and default mode networks) showed the best performance. The default mode network (Network 7) is similar in size to the sensorimotor networks (Networks 1 and 2), whereas Network 6 is much smaller, indicating that predictive success is not merely a consequence of network size (i.e., spatial area and number of vertices). The limbic network also performed poorly, although we note that some regions this network have been associated with signal dropout and related issues so may be less biologically meaningful^17^.

**Figure 5:**
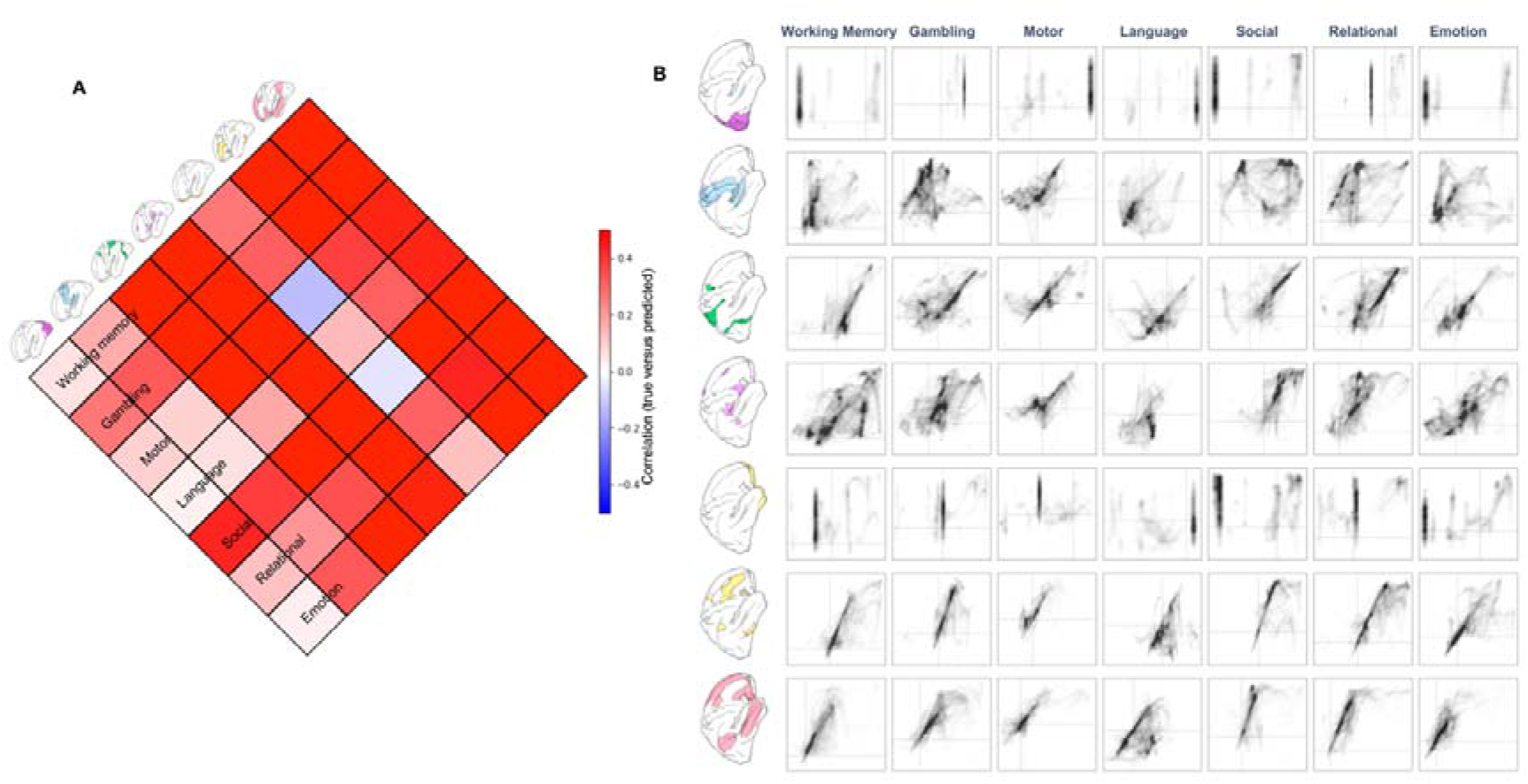
Predicting task activation from masks defined by intrinsic connectivity networks. **A)** The similarity matrix (correlation values) between real and predicted out-of-sample activity profiles; **B)** Scatter plots of the out-of-sample predictions compared to activity for each network and each task.

To assess whether these results arise because of the specific spatial location of each network on the cortical surface, we repeated analyses following spatial or “spin” permutation of each mask on the sphere ^18,19^ and using the location of the rotated masks to make predictions. We found similar out-of-sample predictive performance for true networks and for rotated networks (all p-values > 0.05, FDR or Bonferroni correction). This indicates that the superior predictive performance of heteromodal (than unimodal) networks is due to their spatially distributed nature, rather than the specific location of their sub-regions.

So far we have established that increases in brain activity in humans while they perform tasks, as assessed by fMRI, can be predicted based on the spatial distribution of regions that show decreases in activity. Next we explore two alternative accounts of our data that emerge from contemporary views on the validity of fMRI as a tool for mapping human brain activity: (i) the blurring of signals caused by group averaging and (ii) the lack of biological reality in the fMRI signal.

It is often argued that the method of inter-subject averaging blurs regions with distinct functional profiles so that the time series lose their biologically meaning. To address this possibility we repeated the spatial prediction analysis for three tasks from 10 heavily sampled individual participants from the Midnight Scan Club^20^. Similarly to the group analysis, for each individual a mask formed from the lowest quartile of vertices was used to predict individual activity for vertices across the cortex. As with the group analysis, the predicted activity was generally similar to real activity, both at the individual level and when averaging the results of individualized predictions (Figure 6A and B). We also repeated the pairwise comparisons of different tasks (Figure 4, top) at the individual level, by predicting from the mask of the intersection of lowest quartile of vertices for each pair of tasks. As with the HCP data, we observed greater out-of-sample prediction for true tasks than the alternative tasks (Figure 6C). For 7/10 subjects, prediction was superior for true tasks for all 6 task pairs; for the remaining three subjects, prediction for the true task was superior for 5/6 task pairs. In other words, prediction was superior for the true task for 57/60 task pairs across individual subjects (Figure 6D). This analysis rules out the possibility that our ability to predict increases in brain activity based on the spatial configuration of task-negative activity is an artifact of group averaging.

**Figure 6:**
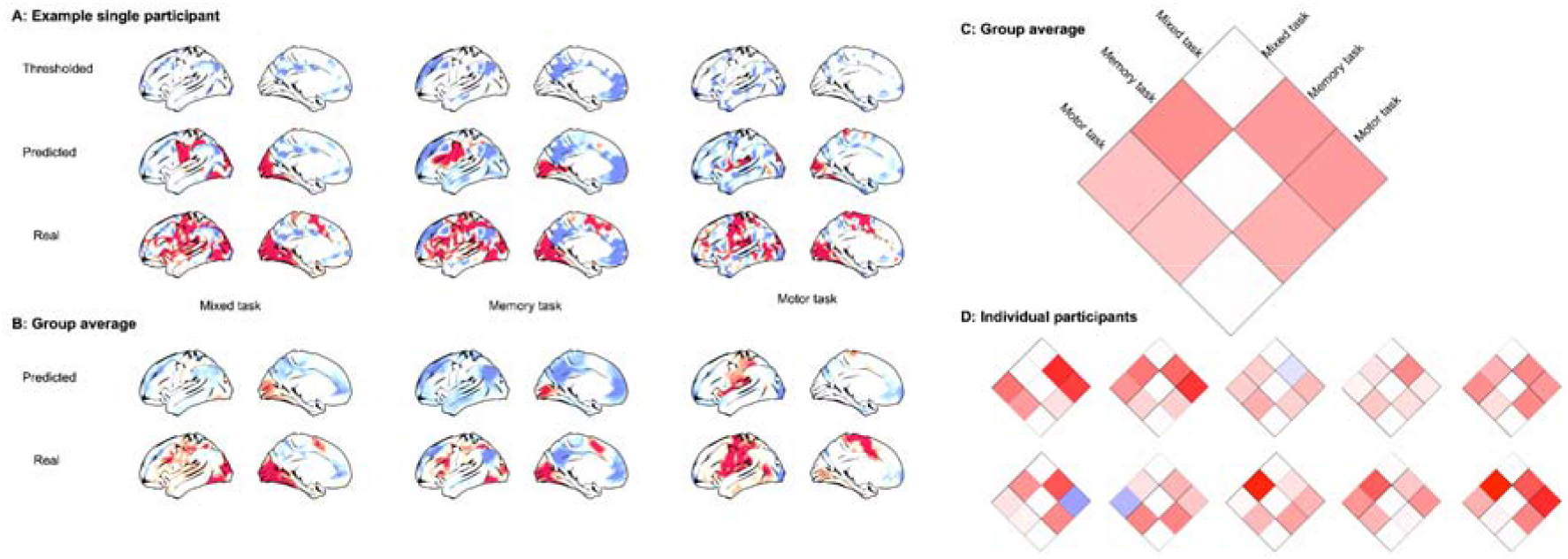
Individual participant analyses: A) The thresholded activity pattern, corresponding prediction and underlying real activity pattern for the three tasks for an illustrative participant projected on an average surface. B) Averaging the individualized predictions and real activity across participants. C) Out-of-sample pairwise similarity (correlation) comparing each task (i.e., the predicted compared to true activity for one task compared to the other tasks). D) The same as C but for each individual participant.

Until now we have considered BOLD responses calculated from human fMRI; however, fMRI is often argued to lack the biological reality that are possible with more direct metrics of neural functions, such as intercranial recordings or calcium imaging. Since our analysis depends on the topography of the cortex, recordings of single regions are unsuitable to test our spatial hypothesis. Instead, we investigated whether this approach generalizes to a more direct metric of neural activity: calcium imaging recordings from visual cortex while a mouse watched a movie (Figure 7). For individual time points, we used the top 25% of pixels to form a mask and used spatial prediction to make out-of-sample predictions (Kriging on a 2D lattice covering the visual cortex). Note, that unlike the human fMRI analyses which were calculated on contrast maps, this analysis used time series data, to predict task-related reductions or increases in activity over time. As with the fMRI data, we observed that long-distance out-of-sample spatial predictions of the pattern of deactivating pixels could be calculated based only on the spatial pattern within the extreme of the distribution. Specifically, we observed positive correlations between real and predicted out-of-sample pixels for 420/540 (78%) time-points, and a mean Spearman correlation of *ρ* = 0.62. This result held even when restricting the out-of-sample prediction to a minimum distance of 25 pixels from the input mask (332/540, 61%, *ρ* = 0.31) (Figure 7D). Our final analysis, therefore, shows that patterns of activity increases in brain activity can be estimated based on geometry from measures of brain activity that are traditionally seen as more biologically plausible than fMRI.

**Figure 7:**
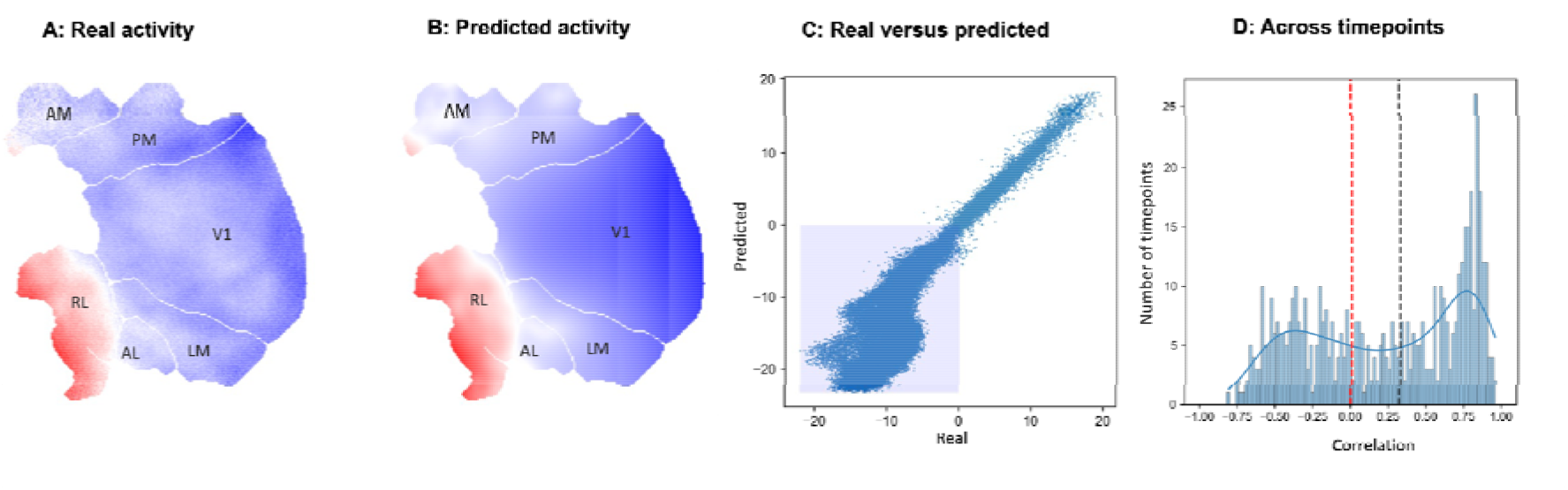
Spatial prediction in a mouse. A: Real activity for a single time point. Labels are: primary visual cortex (V1), lateromedial area (LM), anterolateral area (AL), rostrolateral area (RL), anteromedial area (AM), and posteromedial area (PM). B: Predicted activity from a mask of the top 25% of pixels. C: Real versus predicted activity in out-of-mask pixels (shaded area are negative values - the mask was defined by extreme positive values). D: Top right, distribution of Spearman rank correlations between out-of-sample (>25 pixel distance) real and predicted activity across time points. The black dashed line is the mean correlation.

## Discussion

In this study, we established that the spatial configuration of both increases and decreases in cortical task activity are the result of a common topographic principle. Specifically, across multiple tasks – and for both group averaged and individual data– we found that the spatial organisation of extreme task-negative activity can be used to predict task-positive activity patterns (and vice versa). We could predict not just the spatial location of task-positive activity, but the magnitude of task-positive activity values in a task-specific way. Further, by using intrinsic connectivity networks at rest, we show that networks with different spatial characteristics differ for predicting cortex-wide activity and that this likely reflects the networks’ spatial distribution, rather than its spatial location. Finally, we show in calcium imaging data across mouse visual cortex, that these spatial dependencies are not only restricted to humans or to BOLD functional MRI data.

Our findings indicate that task-positive and task-negative patterns of brain activity, which may be associated with different cognitive processes, are spatially linked and are so likely the result of set of topographical principles that influence how neural processes unfold over space and time. Our results, therefore, can be considered as a spatial analogue of the finding that some brain networks are temporally anti-correlated^21^; our analysis shows that observed activity patterns are such that neural activity is also anti-correlated over space. These spatial constraints are in many ways not surprising given the geometric organisation underlying both the distribution of white matter connections and the preponderance of local connectivity across the cortex^22^. Moreover, they are also predicted by approaches that emphasise how topographic features influence the function of specific brain networks (for example, the default mode network^11^, or the general topography of observed brain activity based on descriptions of its structure^12^.

From a cognitive perspective, one puzzling feature of the task-positive and task-negative features of brain activity is the broad range of situations in which it is important^3^. For example, the multiple demand network, as the name implies, shows a pattern of increased activity across a range of different tasks each of which differ in superficial details (for example they may differ on the type of stimulus, the rate of stimulus presentation and features of task structure). Likewise, regions in the default mode network, such as the posterior cingulate cortex, show reductions in activity in many of the same sorts of tasks that activate during the multiple demand network, and greater activity in situations in tasks that all share a reliance on memory (e.g. semantic knowledge, episodic memory or social cognition)^2,23^. At a cortical level, therefore, the task-positive / task-negative axis of brain function describes spatial patterns of brain activity that occur across a range of specific situations and that may share abstract cognitive features (such as a role of executive control), but also differ in superficial features of the situation in which they are observed (e.g., the Stroop task is not the same as a working memory task). Although the generality of the increases and decreases in brain activity are now well documented, they beg a specific question: What features of cognition are encoded by features of brain activity that are common to tasks that differ in the specific features of cognition and behaviour that they depend upon? In this regard our findings are important because they explain that this persistent spatial motif may occur across many superficially different cognitive settings because of the impact of one or more topographical features that shape how brain activity varies across the cortex. In other words if at least a part of the brain activity pattern we observe during tasks is a result of spatial phenomena which are derived from geometric constraints, then consistent patterns of deactivation and activation should be expected in tasks which may be different in specific features of cognition or behaviour. This is analogous to the way that a small set of common geological processes can explain basic features of landscape of mountains ranges even if they occur in different continents^24^.

Altogether our analysis establishes intimate links between the spatial distribution of increases and decreases in brain activity that can be parsimoniously explained by assuming a set of common topographical principles govern both positive and negative changes in brain activity observed during tasks. Nonetheless, our analysis leaves open several important questions. For example, there are likely to be multiple theoretical approaches that can account for our data. Recently, Pang and colleagues^12^ argued that the neural activity observed during fMRI can be explained by a process in which neural activity is an emergent property of the mechanisms which shape the cortex, arguing that observed brain activity is the result of resonance of spatial features of brain organisation that can be explained by neural field theory. It is possible, therefore, that spatial phenomenon described in our analysis may be explicable in similar terms. Methodologically the spatial regression method used here, Kriging, is a powerful approach applied to a single input brain map, without prior training on a separate dataset (as is common in supervised machine learning). While we only applied this method to functional imaging data, including human fMRI and mouse calcium imaging, this approach can be applied to a wider range of cortical maps, across imaging modalities and species. For example, a relevant candidate for future work are maps of human anatomical, electrophysiological and genetic organisation^25^. Future application of these and related methods (e.g.^26^) to a wider range of maps is likely to further enhance our understanding of the spatial relationships between distributed cortical systems.

Finally, our evidence that geometric constraints play an important role in the observed patterns of brain activity during tasks have important implications for interpreting the links between brain activity, on the one hand, and cognition and behaviour on the other. For example, it is often standard practice to make inferences about a region’s function based on observed increases in functional activity within a specific task. Our data suggests that these inferences could also take into account regions that show reductions in activity, since in many situations the increases in activity contain information about the task context that are also contained in the pattern of reductions. More generally, recent proposals about reservoir computing as a model of cortical information processing ^23^ provide a potential explanation for why both increases and decreases in brain activity across the cortex may be an important feature of the brain basis of cognition. Spatiotemporal dynamics are likely to be relevant when there is a need to balance activation and deactivation either through active homeostatic processes^27,28^ or through careful tuning into a critical regime^29^; this would naturally give rise to task-evoked deactivation patterns as a large-scale spatial homeostatic process^30^. More generally, our findings show that fundamental functional roles of cortex (especially at a meso- and macro-scopic scale) are poorly understood; future work will need to more directly consider the implications of the cortex as a physically embedded spatial system.

## Methods

### Data

#### Group average task data

We used data from the seven group-average task maps from the Human Connectome Project’s Young Adult 1000 participant release. A single contrast map for each task was used (see code for full details). Details of the pre-processing pipeline and creation of the group average maps can be found at^15^. The *xyz*-coordinates for the cortical data were extracted from both the 32k Multimodal Surface Matching^31^ mid-thickness projection (for display purposes), and from the spherical projection (for spatial regression and prediction).

#### Individual participant data

We used individual task contrast maps from the Midnight Scan Club dataset^20^. We used the processed data from the ten participants’ contrast maps for three tasks (Mixed, Memory and Motor), resampled into 32k vertex atlas; full details of the preprocessing are available at OpenNeuro ds000224.

#### Calcium-imaging data

Mouse calcium imaging data was taken from^32^. We used the spatio-temporal data from a single mouse while watching a movie, consisting of 400×400 pixel images (with a pixel size of 0.01mm^2^) acquired at 540 time-points. Details of data acquisition and preprocessing are available at^32^.

#### Spatial prediction with Lattice Kriging

Lattice Kriging (LatticeKrig^14^) is a computationally efficient method for performing Kriging (a form of Gaussian process regression) to large datasets with 10,000s of spatial data points. It involves building a multi-resolution, compactly supported set of basis functions on a regular lattice covering the spatial domain, approximating the covariance structure of the data and allowing for spatial predictions.

##### Human data

For human data, we use the LatticeKrig implementation with spherical geometry (using the spherical projection of the FSLR 32k cortical atlas). We use the default parameters from the LatticeKrig example on the sphere, with three-levels of spatial resolution of basis functions.

The same analysis approach was taken for both group and individual task functional MRI datasets. Vertex values for each contrast map were sorted (after removing the medial wall) and the bottom 25% of vertices were used to create a mask (these were negative values, corresponding to task-negative evoked responses). The spherical coordinates of vertices (calculated from their 3D-coordinates on the sphere) in the mask, and the corresponding task contrast values, were entered into a Lattice Kriging model and used to predict all out-of-sample vertices (i.e., vertices outside of the mask). Subsequent analyses compared prediction on all out-of-sample vertices to the corresponding true values, as well as only the subset of out-of-sample vertices that had positive task-evoked responses.

##### Task-specificity of spatial predictions

To assess whether spatial prediction is task-specific, each task was compared pairwise to each other task (e.g., “Task 1” and “Task 2” below). A conjunction mask was created based on the subset of vertices that were in the bottom 25% for both tasks. To assess prediction, a mask of out-of-sample vertices was created for the remaining 75% of vertex values for Task 1. Vertex values (and their locations) within the conjunction mask were then used to predict out-of-sample vertices based separately on Task 1 and Task 2 values. The predictions from both tasks on out-of-sample vertices were then compared to the true values for Task 1, to assess whether predictive performance was higher for the matching task (Task 1) than for the alternative task (Task 2). This created a non-symmetric out-of-sample pairwise similarity matrix between tasks. Two additional restricted out-of-sample masks were created, as a stricter test of predictive performance: 1) only out-of-sample vertices that also had task-positive values (for Task 1); and, 2) only out-of-sample vertices that were also at distance greater than 0.1 radians from predictor (i.e. input) vertices.

##### Effect of intrinsic network architecture on spatial prediction

A canonical seven-network decomposition of the cortex^16^ was used to create masks of vertices. Task contrast values for all vertices (both positive and negative) within each mask were used to predict vertices outside the mask for each of the seven tasks. To assess whether the spatial location of networks influenced predictive performance, the location/orientation of the networks on the brain was shuffled 500 times using random “spin” rotations of the spherical projection of the cortical surface^18,33^. These were then used to spatially predict task contrast values for out-of-mask vertices; this resulted in a null distribution of predictive performance from the rotated masks^19^.

##### Mouse data

A separate lattice kriging model was performed for each of the 540 time points for the calcium imaging data from a single mouse watching a movie. Given that this is spatio-temporal data rather than a contrast map, there is no equivalent to task positive/task negative pixels. Instead, for each time point, from pixels with a non-zero value, the top 25% of pixels were used to create a mask and the corresponding pixel values and all pixel locations were used to predict the response for: i) out-of-mask pixels; ii) out-of-mask pixels at a minimum distance of 25 pixels from any pixel within the input mask.

For the mouse data, the cortical surface (composed of 57867 pixels) is imaged as a 2-D plane. Therefore, a 2-D regular lattice was used for Kriging, with multi-resolution basis functions with three levels. See code for full details.

Performance was assessed by comparing Fisher transformed correlation values between predicted and real values for each time point. Both t-tests and autoregressive models were used to assess whether the average correlation was greater than chance across time points.

Code to repeat the analyses is openly available at: https://github.com/ActiveNeuroImaging/SpatialAnticorrelations

---

**Supplementary figure 1:**
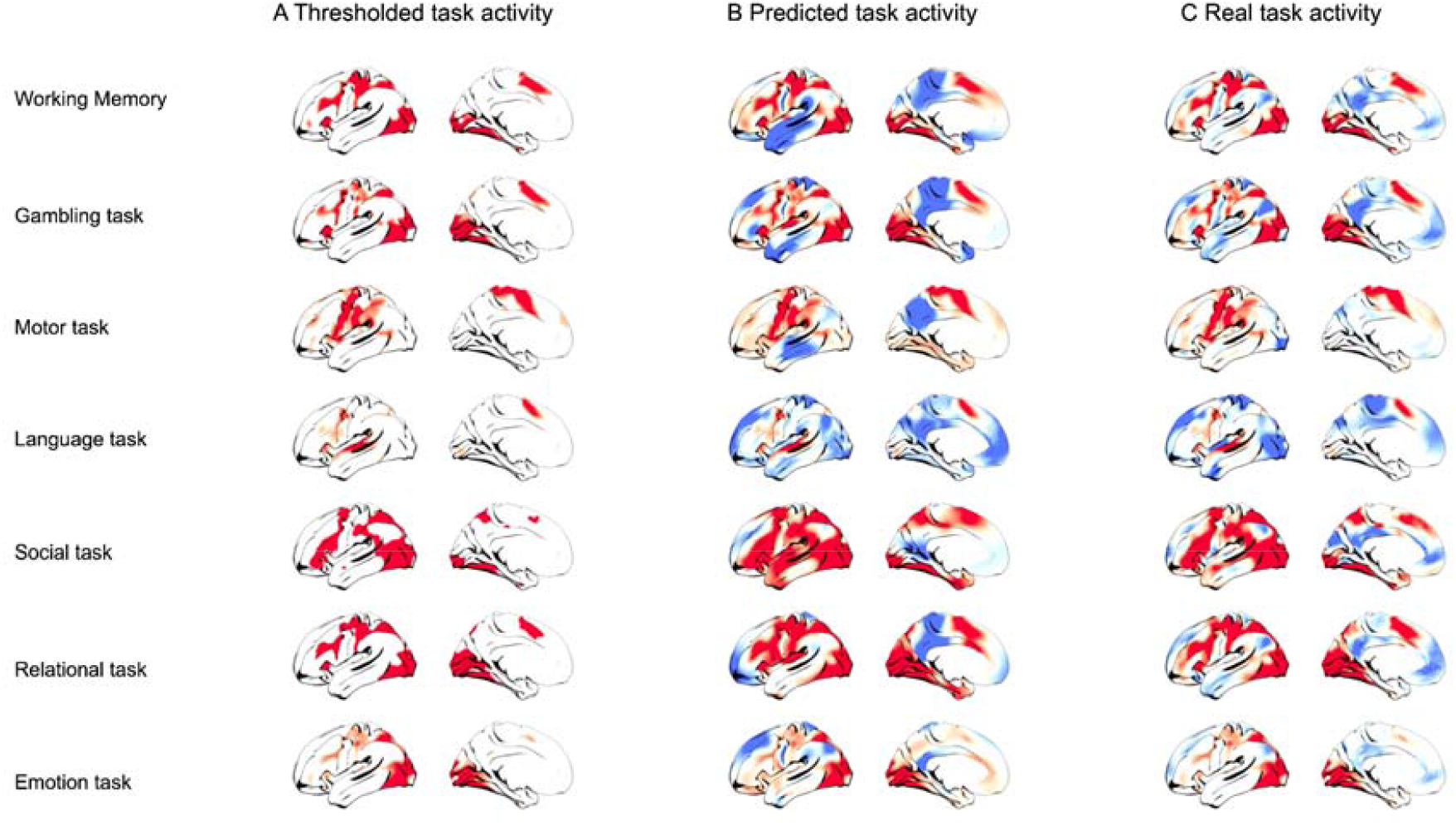
Spatial prediction from task-positive activity patterns. **A**) Vertices with the highest (> 25%) group-average task activation were used to form a predictor mask.**B**) Spatial regression was used to predict task activity at all remaining vertices, and compared to **C)** the true pattern of activity across all vertices.

